# DOUBLER: Unified Representation Learning of Biological Entities and Documents for Predicting Protein–Disease Relationships

**DOI:** 10.1101/2020.10.27.357202

**Authors:** Timo Sztyler, Brandon Malone

## Abstract

**Motivation:** We propose a system that learns consistent representations of biological entities, such as proteins and diseases, based on a knowledge graph and additional data modalities, like structured annotations and free text describing the entities. In contrast to similar approaches, we explicitly incorporate the consistency of the representations into the learning process. In particular, we use these representations to identify novel proteins associated with diseases; these novel relationships could be used to prioritize protein targets for new drugs.

**Results:** We show that our approach outperforms state-of-the-art link prediction algorithms for predicting unknown protein–disease associations. Detailed analysis demonstrates that our approach is most beneficial when additional data modalities, such as free text, are informative.

**Availability:** Code and data are available at: https://github.com/nle-sztyler/research-doubler

*Contact:*

## 1 Introduction

Biological processes within the human body are the result of complex interactions among various molecules, such as DNA, RNA, proteins, metabolites, enzymes, and more. Disease phenotypes result from the disruption of these processes; likewise, effective treatments often aim to restore these processes to their normal, healthy state. Thus, it is imperative to accurately and thoroughly characterize the relationships among the molecules, diseases, and other entities involved in these processes.

Traditional approaches for characterizing these relationships entail expensive and time-consuming biological wet lab experiments. Due to the cost of these experiments, existing datasets are highly incomplete (Härtner *et al.*, 2018; Madeddu *et al.*, 2019). Thus, while this experimental approach remains the gold standard for determining their truth, it is important to devise more cost-effective methods to characterize these relationships.

Increasing evidence demonstrates that artificial intelligence (AI) methods are also very effective for identifying novel relationships (Ghiassian *et al.*, 2015; Caldera *et al.*, 2017; Härtner *et al.*, 2018). Of course, high-confidence relationships predicted by the AI methods can (and should) still be validated with traditional wet lab approaches.

In this paper, we present an approach for learning novel relationships among the various biological entities. We adopt a network-based approach in which each entity is represented as a node in a graph, and each relationship is represented as an edge between the respective entities; different relationship types are represented with different edge types. We refer to this network as a knowledge graph (KG). Learning novel relationships is then equivalent to predicting “missing” edges from the KG (Sharma *et al.*, 2015; Caldera *et al.*, 2017; Vlaic *et al.*, 2018).

Similar network-based approaches have been proposed to address this problem before (Caldera *et al.*, 2017; Madeddu *et al.*, 2019; Malone *et al.*, 2018; Nováček and Mohamed, 2020). The existing approaches can be grouped into three categories (Agrawal *et al.*, 2018): neighborhood-based, diffusion-based, and representation-based. These rely on different topological properties of the network, such as motifs, communities, hubs, clusters, node centrality, shortest paths, and others to predict new edges. Further, some methods also account for biological features of the nodes. For example, proteins may be annotated with their functions (Asif *et al.*, 2018), text from academic papers in which they are described (Zhou and Fu, 2018), or patient sequencing data (Luo *et al.*, 2017); we refer to this additional information as a *document* for the respective entity.

The representation-based approaches are the most similar to ours. These approaches use neural networks to learn *embeddings*, or real-valued numeric vectors, such that entities which are close in the graph and have similar relationships also have similar embeddings. A machine learning model is then trained to predict novel relationships based on the embeddings.

Currently, the proposed approaches do not jointly account for the KG structure and the associated documents in a consistent manner (Agrawal *et al.*, 2018; Vlaic *et al.*, 2018; Ata *et al.*, 2018). In particular, they do not share information between the KG structure and documents about the entities in a unified manner.

We address this shortcoming with a novel neural network learning strategy, DOUBLER (**DO**cuments **U**nified with **B**iological networks for **LE**arning **R**epresentations), which combines the KG structure with the documents in a unified framework in which we force both embeddings to be similar. The main advantage compared to the obvious approach (learning independent embeddings and concatenating them) is that this “pulls together” the embeddings of entities which are distant in the KG but have similar documents; likewise, it “pushes apart” embeddings of genes that are close in the KG but have very different documents. Thus, novel relationships are predicted using a unified view of the KG and documents.

In order to demonstrate the efficacy of our approach, we construct a KG consisting of proteins and diseases. We then compare our approach to existing state-of-the-art bioinformatics and machine learning algorithms for predicting novel protein– disease associations. Empirically, we show that our approach outperforms the competing methods. More detailed analysis demonstrates that our approach is particularly beneficial for cases in which the documents for proteins are informative.

In Section 2, we describe the datasets used to construct the disease–gene network, while Section 3 presents our new approach for learning consistent representations based on documents and the network. The results and discussion are given in Section 4, and Section 5 concludes the paper.

## 2 Materials

DOUBLER requires two data sources: (1) the knowledge graph (KG) and (2) the documents describing entities in the KG.

### 2.1 Biomedical Knowledge Graph

A knowledge graph *G* = (*V, E*) consists of a set of nodes *V* and a set of edges *E*. Each node in the KG corresponds to one entity, such as a protein or disease. The edges characterize the known relationships between these entities.

As an example of constructing the knowledge graph *G* in order to demonstrate and evaluate the proposed approach, we extracted entities and relationships from the following sources.

- `disease - associated_with - protein` DisGeNET: Lo Surdo *et al.* (2017)
- `protein - interacts_with - protein` STRING: Szklarczyk *et al.* (2018)

Additionally, we created `disease - similar_to - disease` relationships using an existing approach (Gottlieb *et al.*, 2011). Further, some data sources provided confidence values for the relationships. We took these into account to filter relationships with a low reliability since those are already predictions.

Our KG *G* includes two types of entities: diseases and proteins. It includes three relationship (edge) types. We use *e*^*t*^ to refer to edge type *t*. In the following, the facts which are encoded by the graph in *G* are represented as a set of triples of the form (h, r, t) where *h* ∈ *V* and *t* ∈ *V* are the head and tail entities and *r* ∈ *E* is a relation type.

As an example, Figure 1 illustrates an excerpt of *G*. It shows that the disease “Asthma” (D1) is associated with “Gene 108” (P2) and “Gene 142” (P1). From a practical point of view, it is expensive to identify novel relations and most of them are still unknown.

**Fig. 1:**
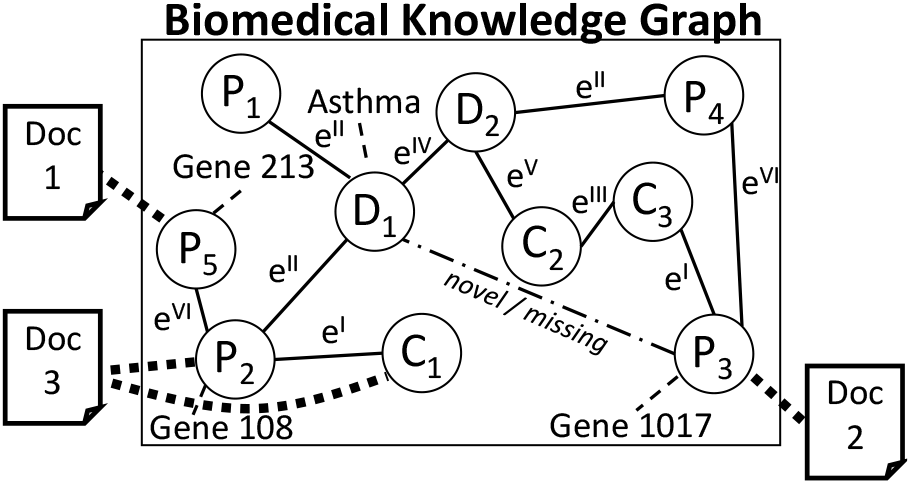
Composition the biomedical knowledge graph (excerpt).

Recently, a variety of similar biomedical KGs have been proposed, such as (Breit *et al.*, 2020) and (Mohamed *et al.*, 2020). Due to the similarity of the source databases, we do not further consider those here; nevertheless, DOUBLER can be directly applied to those KGs, as well.

### 2.2 Mining Biomedical Documents

The documents describe the related entities in the KG in more detail. In particular, we use the following sources and associated types of information.

- Gene Ontology (Gene Ontology Consortium, 2004): protein functions and locations in the cell (structured annotations)
- Human Phenotype Ontology (Köhler *et al.*, 2014): phenotypes associated with diseases (structured annotations)
- MyGene.info (Wu *et al.*, 2013): protein descriptions (free text)
- DisGeNET (Lo Surdo *et al.*, 2017): disease mechanisms and associated mutations (free text)

A common vocabulary is created based on all documents by applying Subword Neural Machine Translation (Sennrich *et al.*, 2016). To reduce vocabulary size, the method breaks non-frequent words into subword units. This is done in an unsupervised manner. As a result, it is possible to encode any (rare) word in subword units. Thus, the vocabulary size is fixed, and we avoid problems from out-of-vocabulary tokens.

Figure 1 illustrates that each document is associated with at least one of the entities from the KG. A document can be associated with more than one entity, and each entity can be associated with more than one document. We assign documents to entities based on existing assignments in the source databases.

Figure 1 illustrates the connection between documents and proteins. More formally, necessary data for DOUBLER is a set of biomedical documents 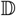, where each 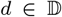 is a multiset of tokens from the vocabulary. Further, 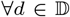, it holds that 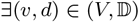 where 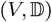 is a set of pairs. We denote this as 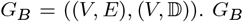 will be used as input for our neural network.

## 3 Methods

We now describe a novel neural network approach for learning to predict novel relationships which combines the knowledge graph (KG) *G_B_* and the related set of biomedical documents 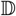 in a unified framework, visualized in Figure 2.

**Fig. 2:**
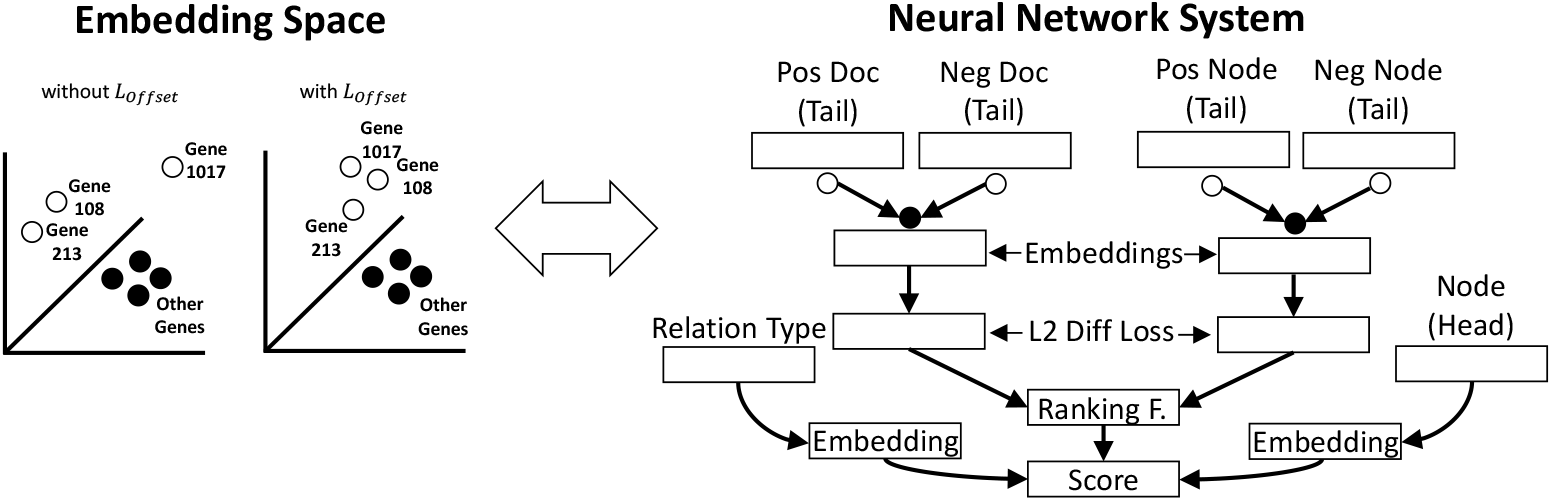
Conceptual connection illustrating how the joined learning of the different modality (right) affects the embedding space (left).

### 3.1 Training

We refer in the following to “proteins” and “diseases” as possible head and tail entities where the corresponding relation is `“associated_with”` (*e*^*d*_*aw*_*p*^). For training the machine learning model, we keep a particular disease (e.g., “Cardiomyopathy”) and the relation *r*_*k*_ (e.g., `“associated_with”`) constant as the head *h* and relation type *r*, respectively, and group the proteins into 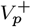 and 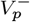 where 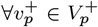 it holds triple in the form of (*h*, `associated_with` 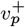) exists in KG. It holds that 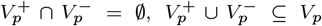 and *V*_*p*_ ⊆ *V*. Analogously, we have 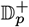 and 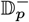 which describe the corresponding biomedical documents of the elements in 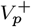 and 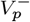. That is, every protein in 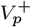 is a correct tail for the triple and is associated with the disease, while every protein in 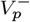 is an incorrect tail and is not known to be associated with the disease. As indicated in Figure 3, we conceptually train our machine learning model with a single sample at a time. That is, we choose 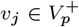 and the corresponding document 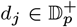. As a first step, this sample is embedded, where the embedding of the nodes *V* and the documents 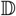 happens independently:

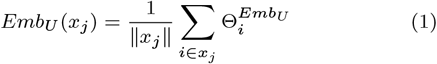

where 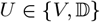, *Emb*_*U*_ is the respective embedding function, 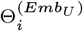 is the embedding of element *i*, *x*_*j*_ is the corresponding element (that is, *v*_*j*_ or *d*_*j*_), *i* = ||*x*_*j*_||. The cardinality for *d*_*j*_ is defined by the number of words in the related documents. The cardinality for *v*_*j*_ is always 1; it is an indicator for the entity in the graph. Second, we apply negative sampling (Kotnis and Nastase, 2018) to randomly select 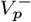 and 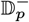, where 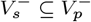 and 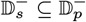. That is, these sets contain incorrect tails for the selected head and relation. Analogously, we embed the set of negative elements as follows:

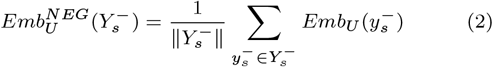

where 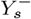 is the set of negative samples (that is, 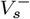 or 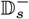). The cardinalities of both 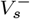 and 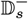 are equal to the number of negative samples. The number of negative samples is a hyperparameter of the method. The embedding of the relation is defined as follows:

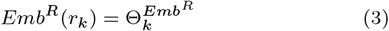

where *k* is the index of a particular relationship type. More concretely, *Emb_V_* is a neural network which gives an embedding for each entity in the KG, while *Emb*^*R*^ is a neural network which gives an embedding for each relationship type. Additionally, 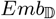 is a neural network which gives an embedding for each token in the vocabulary, and the embedding for a document is simply the average embedding of all of its tokens. 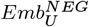 is a simple function that gives the average embedding of all entities that a passed to it.

**Fig. 3:**
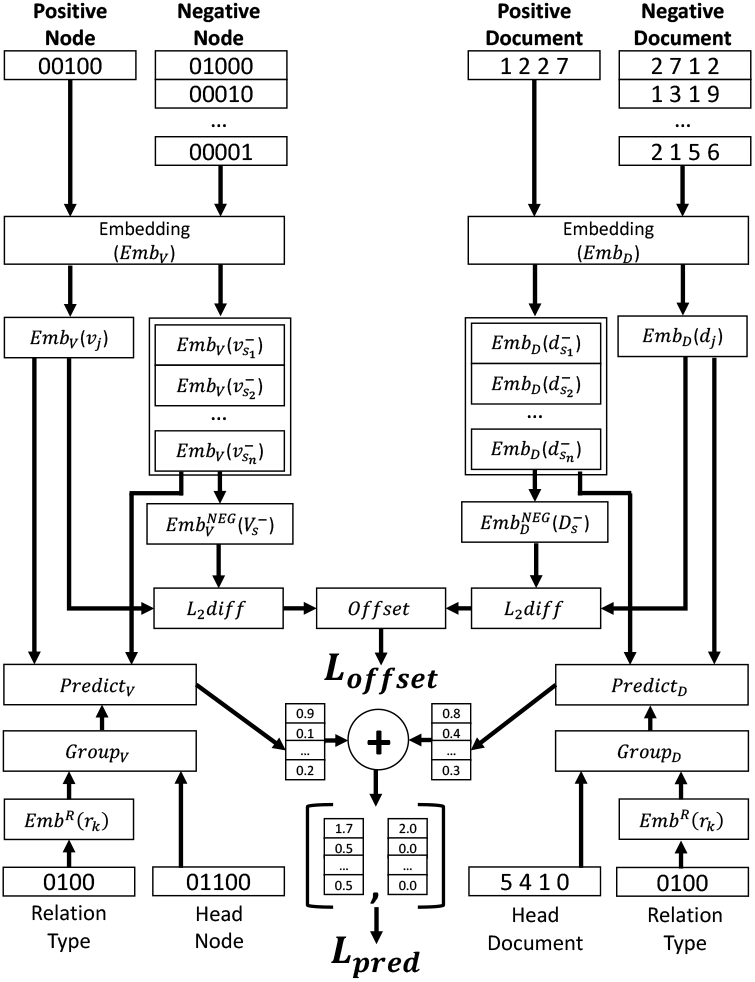
Detailed neural network architecture.

### 3.2 Learning unified embeddings

The neural network calculates the Euclidean distance between the embedding of the positive sample *x*_*j*_ and the center of the negative samples 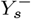:

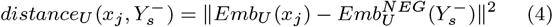

Further, we define *L*_*offset*_ as the distance between *distance*_*V*_ and 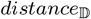:

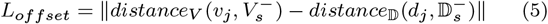

That is, *distance*_*V*_ (*v*_*j*_, 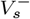) gives the distance between the embedding from the KG for the positive sample and the center of the negative ones, while 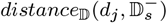 gives the distance between the embedding from the document for the positive sample and the center of the negative ones. In standard approaches, these distances may differ greatly; in such cases, *L*_*offset*_ will be large. In contrast, DOUBLER aims to minimize *L*_*offset*_; hence, the embedding distance between *v*_*j*_ and *d*_*j*_ to the respective negative embedded samples 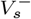 and 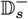 will be minimized. As a result, the model explicitly accounts for similarity in the KG when learning embeddings for the documents and vice versa. That is, the model explicitly unifies the e mbeddings o f the KG s tructure a nd the associated documents. This distinguishes DOUBLER from other methods.

It is important to note that the model adjusts the learned embeddings of the entities to find a b alance b etween the evidence in the KG and the documents. Figure 2 (left) illustrates an example how this can affect the embedding space. The figure s hows t he e mbeddings l earned b y s tandard approaches: Gene 1017 is moderately close in the embedding space to the other Asthma-associated proteins due to the similar documents; however, Gene 1017 is not close to the others in the KG, so it is not clear if it should be associated with Asthma. On the right side of the figure, t hough, w e s ee t he i mpact o f u sing *L*_*offset*_. Due to this unified l earning, t he d ocument e mbeddings inform the KG embeddings. Consequently, in this example, it is now clear that Gene 1017 should be associated with Asthma.

### 3.3 Learning accurate relation predictions

In addition to *L*_*offset*_, the prediction accuracy of the model for predicting relationships is also incorporated into the learning. In particular, it uses the learned weights of the embedding layers (*Emb*_*V*_, 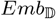, and *Emb*^*R*^) to predict the relationships known to be in the KG. More specifically, recall that we aim to predict the true tail of a triple when the head and relationship type are fixed. T hat i s, w e a re g iven *h* (“Cardiomyopathy” in the example) and *r*_*k*_ (“`associated_with`”), and we wish to predict a correct *t* for the triple (*h, r, t*). We refer to the node and document associated with the head as *v*_*head*_ and *d*_*head*_, respectively.

In contrast to *L*_*offset*_, the prediction accuracy loss *L*_*pred*_ does not average the embeddings of the negative samples (from 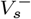 and 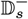) but rather compares all of them to the correct tail (*v*_*j*_ and *d*_*j*_) to measure the predictive performance. In particular, the embeddings of the fixed head and relationship are combined (multiplied element-wise, “·”) to group the nodes in the embedding space by the relation type.

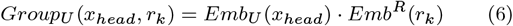

Finally, the score of each candidate tail (that is, *j* as well as each negative sample) is calculated as the dot product of its embedding with *Group*_*U*_ (*x*_*head*_, *r*_*k*_).

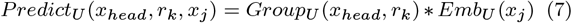

Each score is a measure of how likely the respective entity correctly completes the triple (*h, r,* ?). In the optimal case, the score of the negative samples should be 0 while the true tail should be 1. This step is performed separately for both modalities, and the resulting vectors are combined (added element-wise).

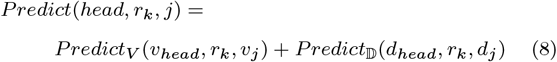

We compute this score for all negative samples and the true tail. Since we consider two modalities, the combined score for the true tail should be 2, while the combined score for the negative samples should still be 0. The resulting scores are considered to compute a binary cross entropy loss.

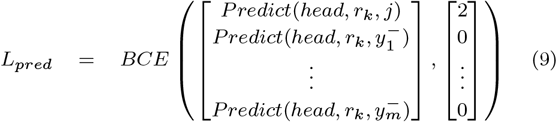

where 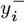 is the *i*^*t*^*h* negative sample and *BCE* is the binary cross-entropy.

Finally, both loss values are combined to measure the performance of the model.

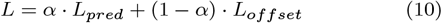

where *α* ∈ [0, 1] is a hyperparameter which controls the tradeoff between accuracy and the unified embeddings. We set *α* = 0.5 in our experiments. Standard backpropagation algorithms are then used to update the parameters of *Emb*_*V*_, *Emb*^*R*^ and 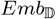 so that the predictions become more accurate while still respecting the unification of the KG and document embeddings; that is, the parameters are updated so that *L* is minimized.

As an example, we refer back to Figure 2. By taking into account the graph structure and the biomedical documents and unifying their embeddings, DOUBLER recognizes that “Gene 1017” (P3) is also associated with Asthma. As a result, we identified a novel relation (*e*^*d*_*aw*_*p*^) which can be investigated in more detail.

### 3.4 Ranking novel relationships

After training the model, the parameters of the embedding functions *Emb*_*V*_, *Emb*^*R*^ and 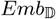 are set such that true triples result in a high *Predict*(·) value, while incorrect triples result in low values. That is, the trained model can answer queries of the following forms: (*h, r,* ?), (*h,* ?*, t*), and (?*, r, t*). Specifically, when given two of the elements, *Predict*(·) is calculated for all possible values of the missing element of the triple. These scores are used to give a ranking from the most-likely (highest-value) to least-likely (lowest) completions of the triple. For cases in which correct completions of the query are known, such rankings can be evaluated using standard information retrieval metrics.

## 4 Results and Discussion

In the following, we describe the conducted experiments and present our results. We compare the performance of DOUBLER with state-of-the-art techniques by predicting disease–protein associations for each disease in our dataset. We present the overall performance but also investigate for which diseases in particular our approach performs better or worse than other techniques. We also characterize the types of diseases for which our method performs well.

### 4.1 Experimental design

#### Dataset

We created a knowledge graph (KG) as described in Section 2.1. We retained only those diseases with at least 10 associated genes. Table 1 shows the characteristics of the knowledge graph.

**Table 1.**
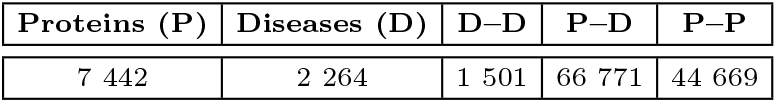
Characteristics of the knowledge graph

#### Methods

Our experiments include evaluation of DOUBLER, as well as five other baselines and state-of-the-art approaches for disease–protein association prediction, including Neural Networks (Grover and Leskovec, 2016), DIAMOnD (Ghiassian *et al.*, 2015), Neighborhood (Navlakha and Kingsford, 2010), RandomWalk (Zhou and Skolnick, 2016), and DistMult (Yang *et al.*, 2014). For DOUBLER and DistMult, we train a single model for all diseases; the remaining methods use disease-specific models.

#### Evaluation metrics

For evaluation, we split the genes associated with each disease into training (80%), validation (10%), and testing (10%) sets. We adopt the commonly used *Recall@100* (*R@100*) metric (Agrawal *et al.*, 2018), evaluated independently for each disease. We further performed 10-fold cross-validation to ensure robustness of the results.

#### Hyperparameter optimization

We did not optimize hyperparameters for DOUBLER. We set them using standard values for similar methods. Specifically, we used a batch size of 1024, generated 100 negative samples for one training sample, and used an embedding dimension of 300 (nodes) or 64 (documents), respectively. The hyperparameters of the other methods were set according to what was reported in the respective papers.

### 4.2 Overall performance

Figure 4 compares the performance of our method to the others. In general, the results illustrate that it is quite challenging to predict valid protein-disease relations; the best-performing methods achieve a mean *R@100* of about 0.4. Further, all of the methods exhibit high variance in performance for different diseases. Nevertheless, DOUBLER outperforms the other methods, demonstrating that incorporating the documents is beneficial.

**Fig. 4:**
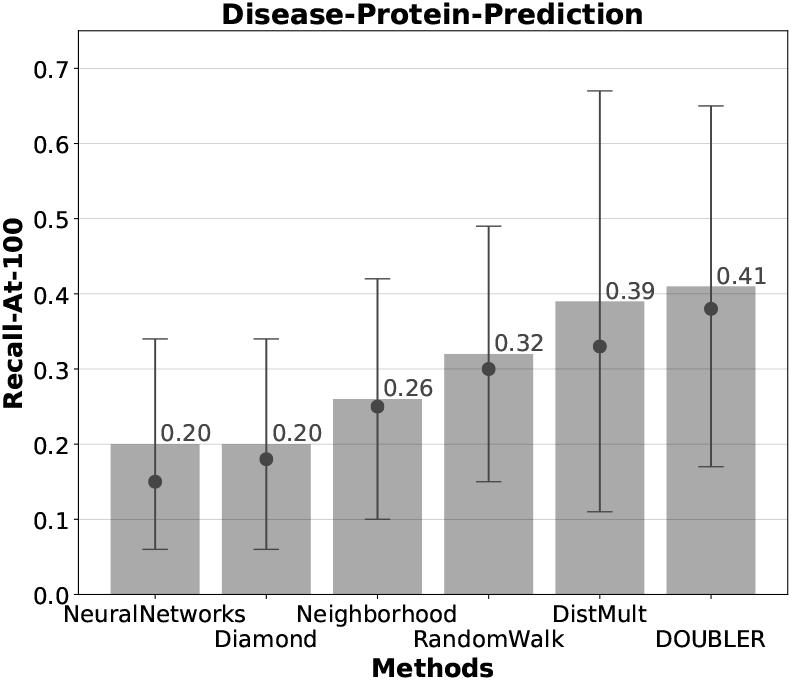
Performance of all methods for predicting disease– protein associations

### 4.3 Convergence behavior

The best performing methods, DOUBLER and DistMult, are both sophisticated neural network-based knowledge graph embedding approaches which can be prone to overfitting. To ensure the stability of the approaches, we monitored their performance on a validation set throughout training. As Figure 5 shows, the *R@100* is quite stable across several different runs with different dataset splits. The figure also highlights that both methods reach their peak performance within a short time, and that DOUBLER continuously performs equal or better than *DistMult*.

**Fig. 5:**
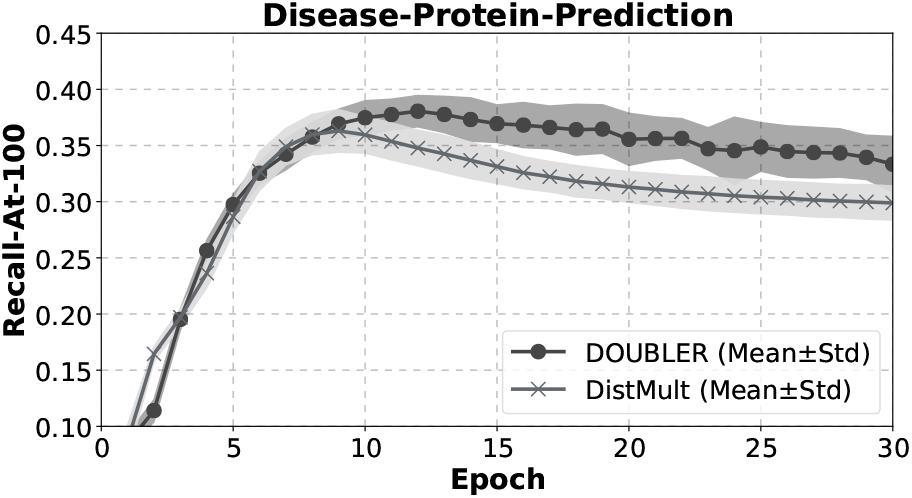
Convergence behavior of DistMult and DOUBLER. The model was evaluated on a held-out validation set. The standard deviation is calculated based on the 10 cross-validation results.

### 4.4 Disease comparison

Comparing the performance of these two methods for each disease individually (Figure 6) shows that, on the one hand, DOUBLER performs similarly or better than DistMult in about 70% of all considered diseases. On the other hand, there are 61 diseases where DistMult performs better.

**Fig. 6:**
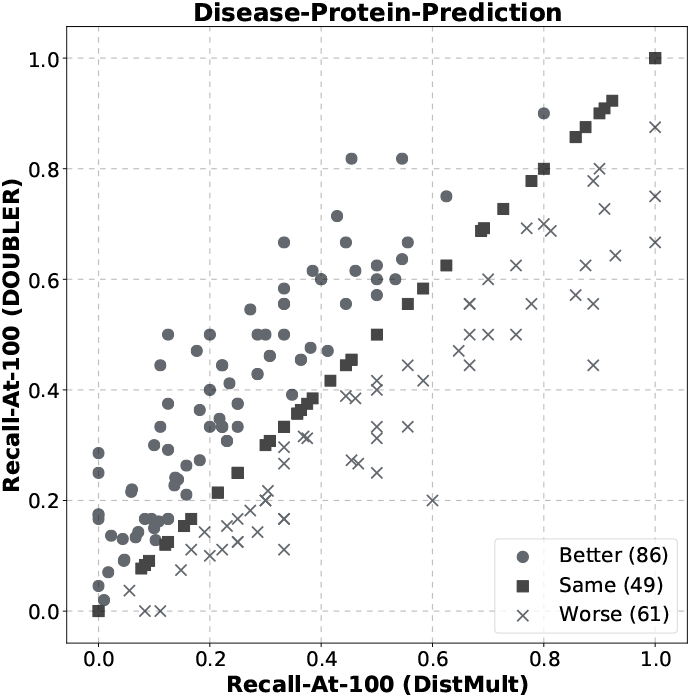
Disease–protein link prediction performance: DistMult vs. DOUBLER. Each point corresponds to a single disease. Circles indicate diseases for which DOUBLER was better, crosses indicate diseases for which DistMult was better, and squares indicate diseases for which performance was equal.

#### Disease and document characteristics

One of the main differences between DOUBLER and DistMult is the use of text documents during learning. In order to better understand the characteristics of the diseases where our approach performs better or worse than DistMult, we extracted those where the gap in *R@100* is > 20% (see Figure 7).

**Fig. 7:**
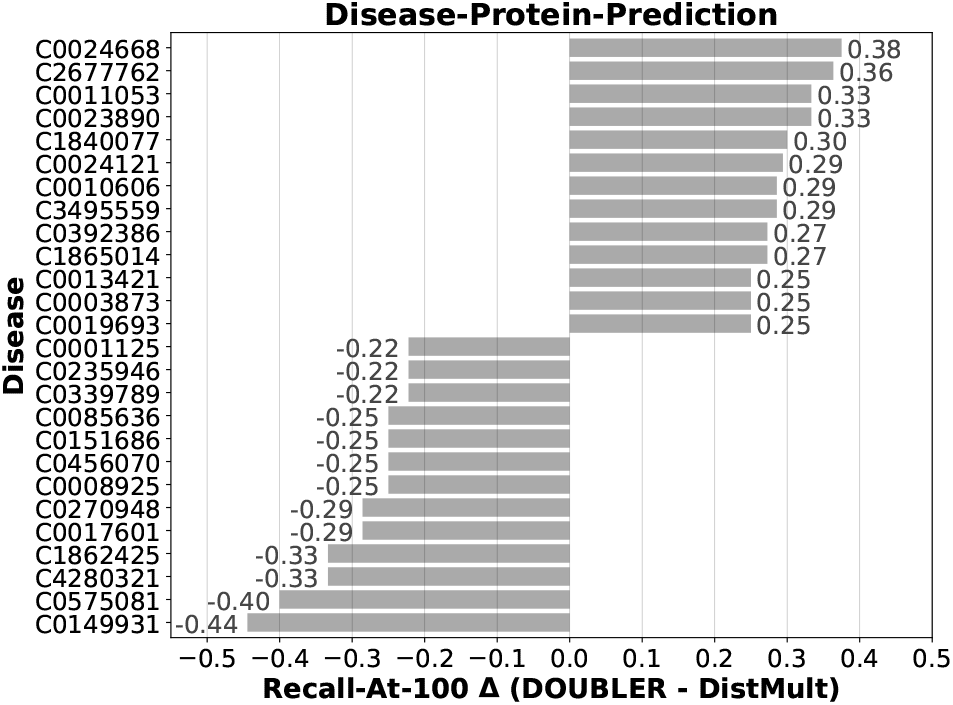
Diseases with the biggest differences in performance for DistMult compared to DOUBLER. Positive values indicate diseases for which DOUBLER was better, while negative values indicate that DistMult was better.

The related protein and document sets show some interesting differences. First, the diseases where DOUBLER performs better tend to have more associated proteins (see Figure 8); consequently, more documents (and, by implication, more training data) are available. Further, Figure 9 shows that the documents related with the diseases that are better recognized by DOUBLER have more unique terms per protein. This suggests that those documents describe the proteins in more detail. Since DOUBLER incorporates the documents into the learning, it leverages these detailed descriptions to train better models.

**Fig. 8:**
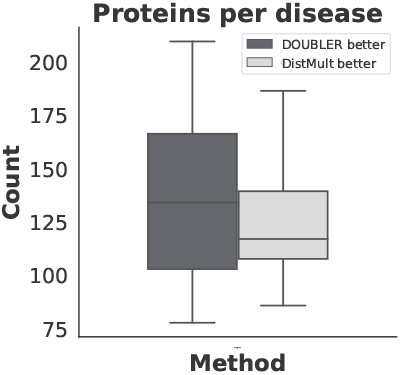
Number of proteins per disease

**Fig. 9:**
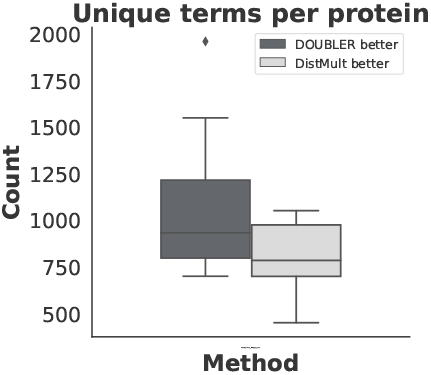
Unique terms per protien.

We next analyzed the individual terms within the documents for which the performance of DistMult or DOUBLER was different. We first considered the inverse document frequency (IDF) of terms in the documents. In general, a high IDF shows that a term occurs only in a few documents, while a low IDF indicates that a term appears in many documents and may not be very informative. Indeed, our analysis (Figure 10) shows that DOUBLER outperforms DistMult when the document terms have a high IDF and are informative about the disease. Finally, we investigated the usage of each term within the documents for each disease (Figure 11). As expected, DOUBLER outperformsDistMult for diseases in which terms are more frequently used in multiple documents for the disease.

**Fig. 10:**
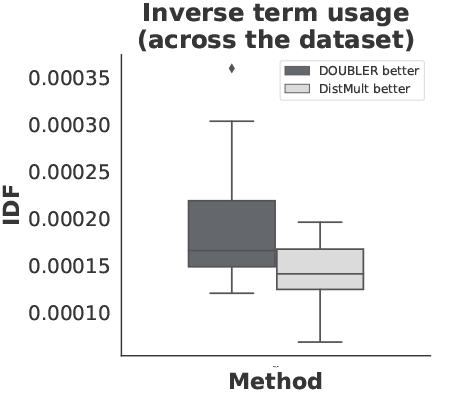
Inverse term usage (across the dataset)

In summary, these results paint a consistent picture in which DOUBLER outperforms DistMult and other state-of-the-art approaches for complex diseases which involve many genes (Figure 8), and also where the individual genes within the disease are consistently described (Figures 9–11). While these results are not surprising considering that a key difference of DOUBLER compared to others is its use of documents, these quantitative and qualitative experiments confirm this expected behavior.

**Fig. 11:**
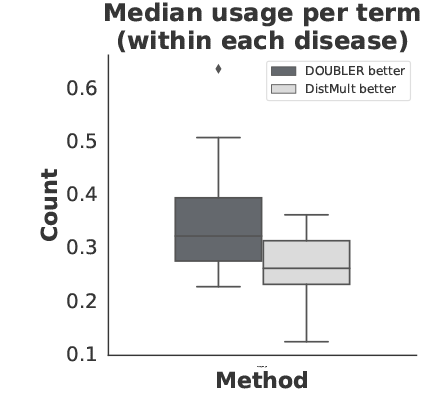
Fig. 11: Median usage per term (within each disease)

## 5 Conclusions and Future Work

In this work, we have presented a general framework for learning representations of biological entities which are consistent among both data attributes and the network structure. In particular, we incorporate a Euclidean distance offset between embeddings based on the knowledge graph structure and the documents associated with entities into the loss function for training a neural network; thus, the model explicitly unifies the embeddings of the KG structure and the associated documents. This distinguishes our approach from other proposed methods. We demonstrated that this approach outperforms competing methods, and more detailed analysis showed it is particularly beneficial when the data attributes are informative. Future work will evaluate the efficacy of our approach for more comprehensive knowledge graphs, such as the recent OpenBioLink (Breit *et al.*, 2020) dataset. Further, it is well known (Kadlec *et al.*, 2017) that hyperparameter tuning can lead to significant performance gains for many methods; thus, future work will also incorporate hyperparameter optimization strategies.

